# The narrow-spectrum anthelmintic oxantel is a potent agonist of a novel acetylcholine receptor subtype in whipworms

**DOI:** 10.1101/2020.09.17.301192

**Authors:** Tina V. A. Hansen, Susanna Cirera, Cédric Neveu, Kirstine Calloe, Dan A. Klaerke, Richard J Martin

**Affiliations:** Department of Veterinary and Animal Sciences, Faculty of Health and Medical Sciences, University of Copenhagen, Dyrlaegevej 100, 1870 Frederiksberg C, Denmark; INRAE, UMR Infectiologie et Santé Publique, Nouzilly, France; Department of Biomedical Sciences, College of Veterinary Medicine, Iowa State University, Ames, IA, USA

## Abstract

In the absence of efficient alternative strategies, the control of parasitic nematodes, impacting human and animal health, mainly relies on the use of broad-spectrum anthelmintic compounds. Unfortunately, most of these drugs have a limited single-dose efficacy against infections caused by the whipworm, *Trichuris*. These infections are of both human and veterinarian importance. However, in contrast to a wide range of parasitic nematode species, the narrow-spectrum anthelmintic oxantel has a high efficacy on *Trichuris spp*. Despite this knowledge, the molecular target(s) of oxantel within *Trichuris* is still unknown. In the distantly related pig roundworm, *Ascaris suum*, oxantel has a small, but significant effect on the recombinant homomeric **N**icotine-sensitive ionotropic acetylcholine receptor (*N*-AChR) made up of five ACR-16 subunits. Therefore, we hypothesized that in whipworms, a putative homolog of an ACR-16 subunit, can form a functional oxantel-sensitive receptor. Using the pig whipworm *T. suis* as a model, we identified and cloned a novel ACR-16-like receptor subunit and successfully expressed the corresponding homomeric channel in *Xenopus laevis* oocytes. Electrophysiological experiments revealed this receptor to have distinctive pharmacological properties with oxantel acting as a full agonist, hence we refer to the receptor as an *O*-AChR subtype. Pyrantel activated this novel *O*-AChR subtype moderately, whereas classic nicotinic agonists surprisingly resulted in only minor responses. We demonstrated that the novel *Tsu-*ACR-16-like receptor is indeed a target for oxantel and is more responsive to oxantel than the ACR-16 receptor from *A. suum*. These finding most likely explain the high sensitivity of whipworms to oxantel, and highlights the importance of the discovery of additional distinct receptor subunit types within *Trichuris* that can be used as valuable screening tools to evaluate the effect of new synthetic or natural anthelmintic compounds.

**Author Summary:** The human whipworm, *Trichuris trichiura*, is an intestinal parasitic nematode infecting approximately 289.6 million people globally, primarily children living in developing countries. Chronic *T. trichiura* infection may cause dysentery, growth stunting and decreased cognitive performance. Whipworm infections are notoriously difficult to control with most available anthelmintics, including those commonly used in mass drug administration programs. Recently performed randomised controlled trials with whipworm-infected humans, have reported superior efficacies of oxantel, a classic, narrow-spectrum anthelmintic, developed for the treatment of *Trichuris* infections. Despite this knowledge, the molecular target(s) of oxantel within the whipworm has not been identified. In this study, we used the whipworm from pigs as a model and identified a receptor, which was explored using the *Xenopus* oocyte expression system. We demonstrated that this receptor is highly responsive to oxantel, and therefore a major target of oxantel within *Trichuris*. In addition, we discovered that this receptor-type is distinctive and only present in the ancient group of parasitic nematodes, Clade I, which also includes the important zoonotic parasite *Trichinella*. Our findings, explain the specific mode of action of oxantel and open the way for additional characterization of similar receptor subtypes in other medically or veterinary important parasitic nematodes of Clade I.

## 1. Introduction

The human whipworm, *Trichuris trichiura*, is a Clade I parasitic nematode [1] and one of the Soil Transmitted Helminths (STHs) that is estimated to infect 289.6 million people globally, primarily those living in the tropics and subtropics [2]. Trichuriasis is rarely fatal, but chronically affects the health and nutritional status of the host [3,4], and is known to be notoriously difficult to treat using current anthelmintic drugs (e.g. albendazole and mebendazole) [5–12]. The extensive use of anthelmintics in livestock has led to widespread anthelmintic resistance (AR) to all the major drug classes [13]. Therefore AR in human parasitic nematodes is a concern where decreased susceptibility to albendazole has already been reported for both *T. trichiura* [14] and the human roundworm, *Ascaris lumbricoides* [15].

Oxantel, is a cholinergic agonist [16], and a *m*-oxyphenol analogue of pyrantel which was developed in 1972 [17] and marketed as a veterinary anthelmintic in 1974 for the treatment of *Trichuris* [18]. Early clinical trials reported oxantel to be effective against *T. trichiura* infections [19,20] and recent studies show that oxantel is superior to single-dose albendazole and mebendazole [21,22], which are currently recommended by the WHO for the control of STHs [23]. Cholinergic agonists [16] exert their effect by paralyzing the worms, which are subsequently killed or expelled from the host [24]. This effect is mediated by nicotinic acetylcholine receptors (nAChRs) [24] that are either heteromeric or homomeric five-subunit ligand-gated ion channels expressed in neuronal, muscle and non-neuronal cell membranes [25,26]. nAChRs of parasitic nematodes have been separated into different pharmacological subtypes based on their sensitivities to a range of cholinergic anthelmintics. Patch-clamp recordings of muscle cells isolated from the pig roundworm *A. suum*, have revealed that their muscle nAChRs are preferentially activated either by levamisole (L), nicotine (N) or bephenium (B), and correspondingly are described as *L*-, *N*-, and *B*- AChRs subtypes [27]. Oxantel is classified as an agonist which is selective for the *N*-AChR subtypes [16]. The *N*-AChR subtypes from the model nematode *Caenorhabditis elegans* and the distantly related pig parasite *A. suum* are homomeric receptors made of the ACR-16 subunits [28,29]. Both of these ACR-16 receptors have a low, but significant sensitivity to oxantel [29,30].

The high sensitivity of *Trichuris* spp. to oxantel has previously been speculated to be due to an nAChR subtype present in *Trichuris* spp. that differs from nAChRs present in other intestinal parasitic nematodes [16]; we hypothesized that a potential homolog of ACR-16 in *Trichuris* could be a target of oxantel within this species.

Here we describe the functional characterization of a novel AChR subtype from the pig whipworm *T. suis* with a high sensitivity to oxantel and distinctive pharmacological properties. This homomeric receptor, referred to as an *O*-AChR subtype, is made of a divergent subunit specific to Clade I nematode species that is only distantly related to ACR-16 from nematode species belonging to other clades. Our results provide new insights about the mode of action of oxantel, its high efficacy on whipworms, and the divergent anthelmintic sensitivity of whipworms.

## 2. Results

### 2.1. Identification of T. suis sequences related to the ACR-16 group

Using the *C. elegans* ACR-16 deduced amino-acid sequences as a query, tBLASTn search against *T. suis*, *T. muris* and *T. trichiura* genomic data available in WormBase-ParaSite (version WBPS14; http://parasite.wormbase.org/) allowed the identification of two distinct hits for each species sharing identities ranging from 46% to 47% with the *C. elegans* sequence. Subsequently, a second tBLASTn search against nematodes genomic data available at the NCBI was performed with the retrieved *Trichuris spp*. sequences. Homologies could be identified with either of the *acr-16* or *acr-19* genes from nematode species representative from the nematoda phylum. Using a panel of representative *C. elegans* nAChRs subunits as reference, a phylogenetic analysis including the *Trichuris* spp. and their putative homologs in the closely related species *Trichinella spiralis* was carried out (Fig. 1). *Trichuris* spp. sequences were found to form two distinct clusters. The first one presented a clear orthologous relationship with ACR-19, the second one clustered apart from the other subunits and belonged to the ACR-16 group [31]. An additional analysis, including other related sequences from nematode species representative from the different clades of the nematoda phylum (Suppl. Fig. 1) further confirmed that Clade I nematode species (including *Trichuris* spp.) possess a divergent group of AChR subunit related to the ACR-16 subunit. Consequently, the corresponding sequences from Clade I nematode species were named ACR-16-like.

**Figure 1.**
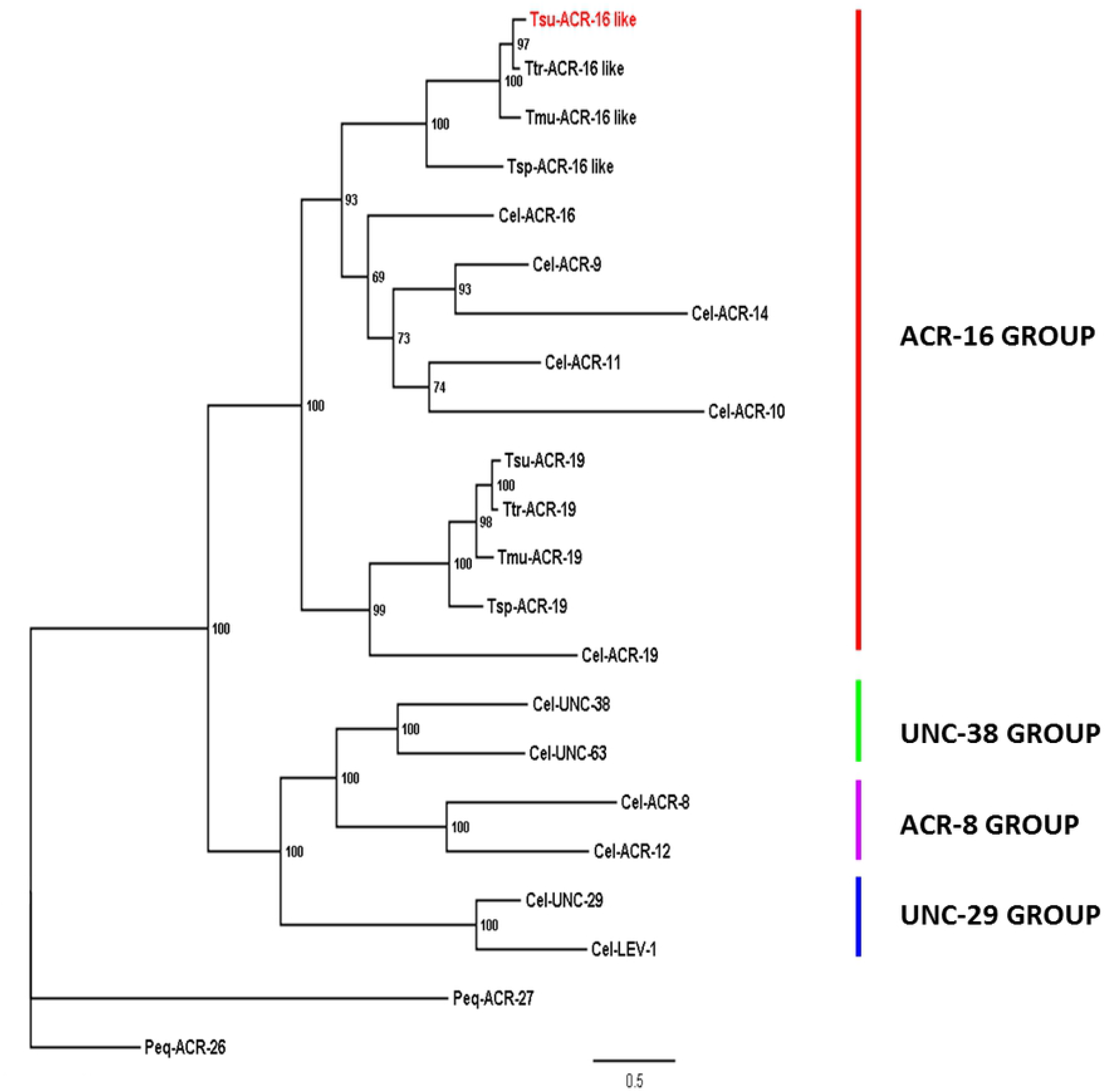
Maximum likelihood tree showing relationships of the ACR-16 related acetylcholine receptor (nAChR) subunits from *Trichuris species,* with other *C. elegans and T. spiralis* nAChR subunits. Tree was built upon an alignment of nAChR subunit deduced amino-acid sequences. The tree was rooted with the *Parascaris equorum* ACR-26 and ACR-27 sequences that are absent from *C. elegans* and clade I nematode species (Courtot *et al.* 2018). Scale bar represents the number of substitutions per site. Boostrap values are indicated on branches. Accession numbers for sequences used in the phylogenetic analysis are provided in the Material and Methods section. *C. elegans* nAChR subunit groups are named as proposed by Mongan *et al.* [31], *Cel*, *Tsu*, *Ttr*, *Tmu*, *Tsp* and *Peq* refer to *Caenorhabditis elegans*, *Trichuris suis*, *Trichuris trichiura*, *Trichuris muris* and *Parascaris equorum,* respectively.

### 2.2 Molecular cloning of the Tsu-acr-16-like coding sequence

In the present study, based on the current knowledge of the original mode of action of oxantel [16,29], we hypothesized that the divergent ACR-16-like from *Trichuris* spp. could represent a preferential target for this narrow-spectrum anthelmintic. Using *T. suis* as a model, we cloned its *acr-16-like* full-length cDNA sequence as a matter of priority and deposited the sequence in GenBank under the accession number MT386096. An alignment of the *Tsu*-ACR-16-like sequence with its closely related counterparts from *T. muris*, *T. trichiura*, *T. spiralis* and *C. elegans* is provided in Fig. 2. The *Tsu-*ACR-16-like subunit was found to share typical features of nAChR subunits including a predicted signal peptide, a Cys-loop motif, four transmembrane regions (TM1-TM4), and the YxCC motif which characterize an α-type nAChR receptor subunit.

**Figure 2.**
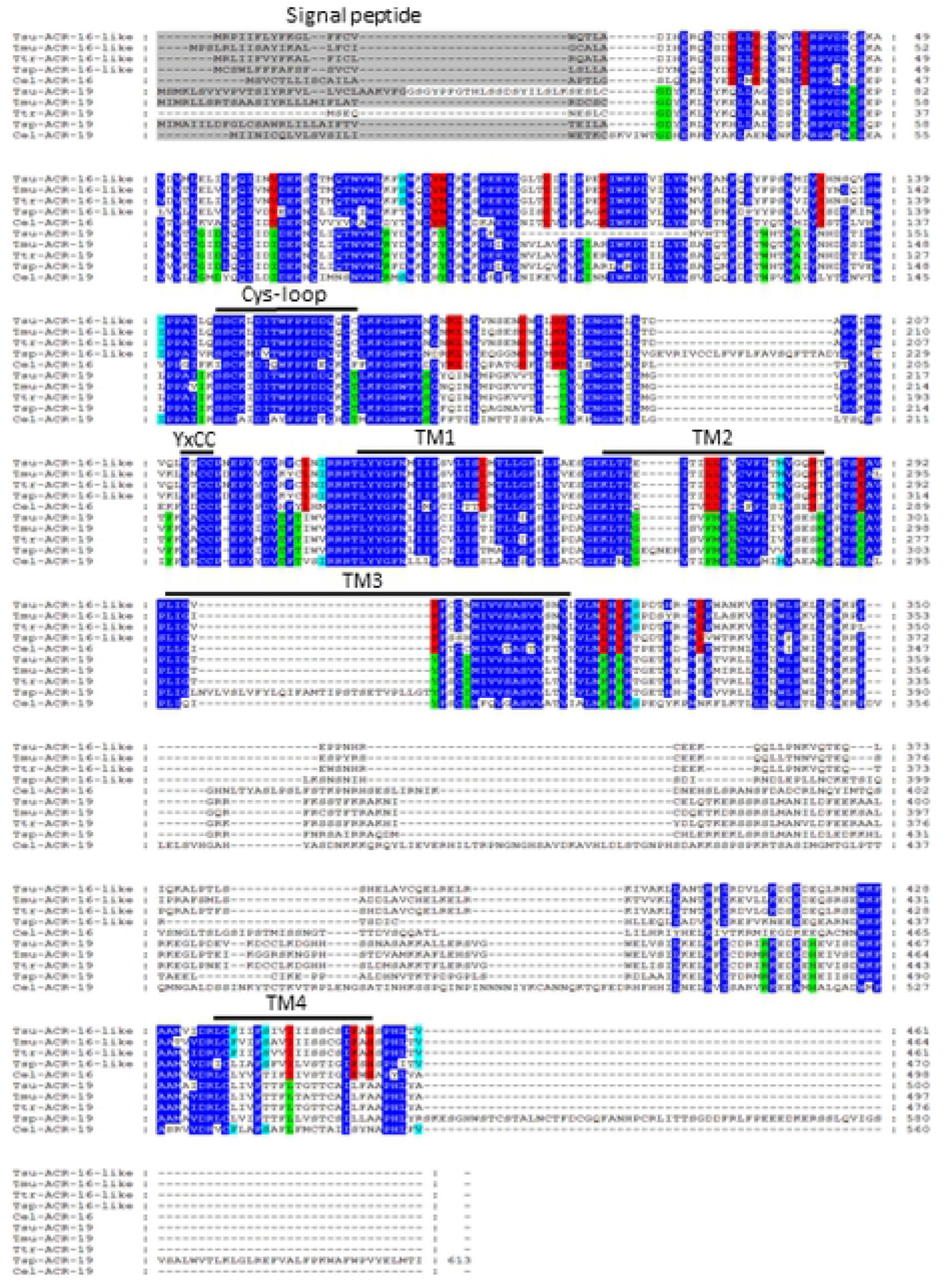
Amino acid alignment of ACR-16-(like) and ACR-19 subunit sequences from the Clade I parasitic nematodes *Trichuris suis*, *T. trichiura*, *T. muris*, *Trichinella spiralis* and *Caenorhabditis elegans*. Predicted signal peptide sequences are shaded in grey, the Cys-loop, the transmembrane regions (TM1-TM4), and the YxCC motif that characterize an α-subunit are indicated above the sequences. Conserved amino acids between ACR-16-(like) and ACR-19 sequences (dark blue), conserved amino acids between all ACR-16-(like) sequences (red), conserved amino acid between all ACR-19 sequences (light green), conserved amino acid between ACR-16-like sequences of Clade I parasitic nematodes and ACR-19 of *C. elegans* (light blue).

### 2.2. Tsu-ACR-16-like subunits form a functional homomeric receptor when co-expressed with the ancillary protein RIC-3 in the Xenopus laevis oocyte

Previous studies have shown that *Cel*-ACR-16 and *Asu-*ACR-16 can form functional homomeric receptors when expressed in *X. laevis* oocytes with the ancillary protein RIC-3 [29,32]. Thus, we explored the requirement of RIC*-*3 for optimizing the putative functional expression of *Tsu-acr-16-like* cRNA in *X. laevis* oocytes. *Xenopus laevis* oocytes were micro injected with *Tsu-acr-16-like* cRNA in combination with *Asu-ric*-3 or *Xle-ric-*3 cRNA. *Tsu-acr-16-like* cRNA or *Asu-ric*-3 cRNA alone as well as non-injected oocytes were used as controls.

The combination of *Tsu-acr-16-like* and *Asu-ric*-3 or *Tsu-acr-16-like* and *Xle-ric-*3 cRNAs led to robust expression of functional receptors responding to 100 μM ACh, which elicited inward currents in the μA range (Fig. 3). The largest currents were measured in oocytes co-injected with *Tsu-acr-16-like* and *Asu-ric*-3 cRNA (mean ± SEM, 7.54 ± 0.74, μA, *n*=32, data not shown); subsequently all other responses were normalized to this. Oocytes co-injection with *Tsu-acr-16-like* and *Xle-ric*-3 cRNA resulted in relatively high current responses (mean ± SEM, 5.0 ± 0.72 μA, *n*=6, data not shown), but oocytes injected with this cRNA combination degraded faster than oocytes injected with *Tsu-acr-16-like* and *Asu-ric*-3 cRNA (data not shown). Non-injected oocytes, or oocytes injected with either *Tsu-acr-16-like* or *Asu-ric*-3 cRNA alone, did not respond to 100 μM ACh, highlighting the need of RIC-3 for the functional expression of the homomeric *Tsu*-ACR-16-like receptor. Representative traces of the inward currents for each injection type, and a bar chart presenting their mean ± SEM normalized values are shown in Fig. 3. Based on these results, all subsequent recordings were performed on oocytes co-injected with *Tsu-acr-16-like* and *Asu-ric*-3 cRNAs.

**Figure 3.**
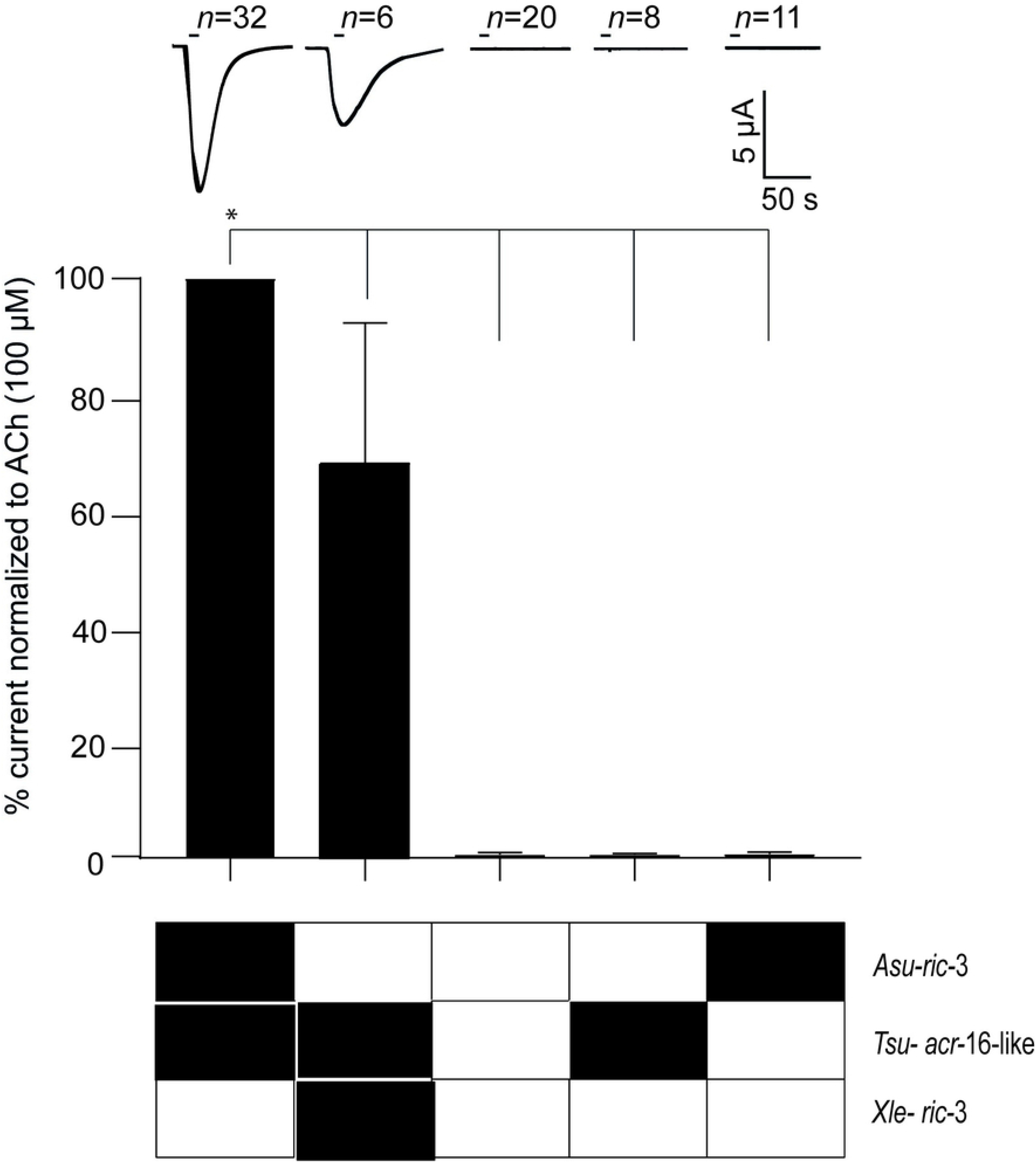
Effect of the ancillary protein Resistance-to-cholinesterase (RIC-3) from *Ascaris suum* (*Asu-*RIC-3) and *Xenopus laevis* (*Xle-*RIC-3) on the functional expression of the ACR-16-like nAChR from *Trichuris suis* (*Tsu-*ACR-16-like receptor). Representative sample traces of inward current in response to 100 μM ACh are shown together with a bar chart presenting the relative currents (mean ± SEM). Oocytes injected with *Tsu-acr-16-like*- or *Asu-ric-3* cRNA alone did not respond to 100 μM ACh. The relative current of oocytes co-injected with *Tsu-acr-16-like* and *Asu-ric*-3 cRNA was significantly higher than oocytes injected with *Tsu-acr-16-like* and *Xle-ric*-3 cRNA. *P* < 0.05; significantly different as indicated, Dunnett’s test.

### 2.3. Oxantel is a potent agonist on the Tsu-ACR-16-like receptor

To explore the effect of oxantel and perform a detailed pharmacological characterization of the *Tsu-* ACR-16-like receptor, we used 4 cholinergic anthelmintics (i.e. oxantel, pyrantel, morantel and levamisole) and 5 nAChR agonists (i.e. epibatidine, nicotine, 3-bromocytisine, DMPP and cytisine). Fig. 4 shows the rank order potency series of these drugs, representative traces of the inward currents induced by each of them, the number of oocytes (*n*) used for each agonist, along with a bar chart presenting the normalized mean ± SEM for each drug group. Oxantel was the most potent agonist on the *Tsu-*ACR-16-like receptor and induced a current response even higher than the control response of 100 μM ACh. Pyrantel also induced a relatively high current response, however, the nicotinic agonists: epibatidine, nicotine, 3-bromocytisine, DMPP and cytisine as well as the cholinergic anthelmintics, morantel and levamisole, were the least potent. The rank order potency series for the agonist drugs on the *Tsu-*ACR-16-like receptor when normalized to control oocytes exposed to 100 μM ACh was: oxantel > ACh >>> pyrantel >>> epibatidine > nicotine ~ 3-bromocytosine ~ DMPP ~ morantel ~ cytosine ~ levamisole. Taken together, these observations provide strong evidence that the *Tsu*-ACR-16-like receptor represents the preferential molecular target for oxantel. Interestingly, when exposed to 100 μM ACh for 1-3 min the *Tsu-*ACR-16-like receptor did not show a fast-desensitization kinetics (Suppl. Fig. 2) which is a distinctive characteristic of the *N*-AChR of nematode such as *Asu*-ACR-16 [29] and the ACR-16 from *Parascaris equorum* (*Peq*-ACR-16) [33].

**Figure 4.**
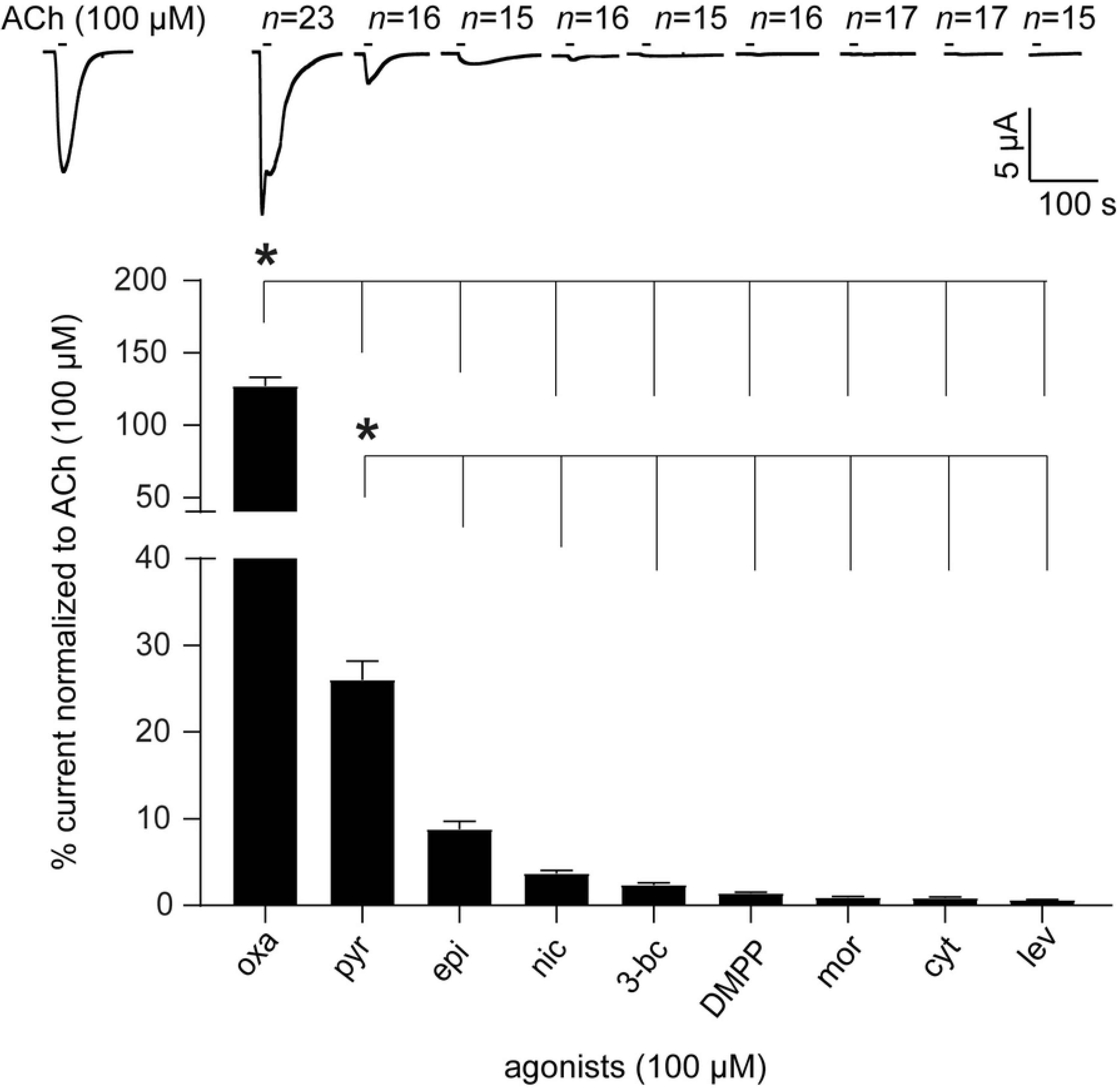
The effect of 9 agonists on *Tsu-*ACR-16-like receptor. Representative sample traces and a bar chart (mean ± SEM) show the rank order potency series of 4 cholinergic anthelmintics: oxantel (oxa), pyrantel (pyr), morantel (mor), levamisole (lev) and 5 nAChR agonists: epibatidine (epi), nicotine (nic), 3-bromocytisine (3-bc), dimethylphenylpiperazinium (DMPP) and cytisine (cyt). *P* < 0.05; significantly different as indicated; Turkey’s multiple comparison test.

### 2.4. Dose-response curve of oxantel and pyrantel

Oxantel and pyrantel have similar chemical structures (Fig. 5A), but their potencies on the *Tsu-*ACR-16-like receptor were found to be significantly different (Fig. 4). We performed a dose-response study on oxantel and pyrantel to determine their *EC*_*50*_ values, their relative maximum current responses, *I*_max,_ and their Hill slopes, *n*_H_. The mean current response of positive control oocytes exposed to 300 μM ACh was used for normalization. Fig. 5B shows representative traces and dose-response relationships of the normalized inward currents (mean ± SEM) induced by different concentrations of oxantel or pyrantel. The *EC*_*50*_ ± SE for oxantel (9.49 ± 1.13 μM) was significantly lower than that of pyrantel (148.5 ± 1.19 μM, *P* < 0.001). The relative maximum current response, *I*_*max*_ (mean ± SE) was significantly larger for oxantel (86.85 ± 4.63%) than pyrantel (29.41 ± 1.95%, *P* = 0.003), whereas no significant difference was found between the Hill slopes, *n*_H_ (mean ± SE), of oxantel (2.51 ± 1.31) and pyrantel (3.13 ± 1.07).

**Figure 5a.**
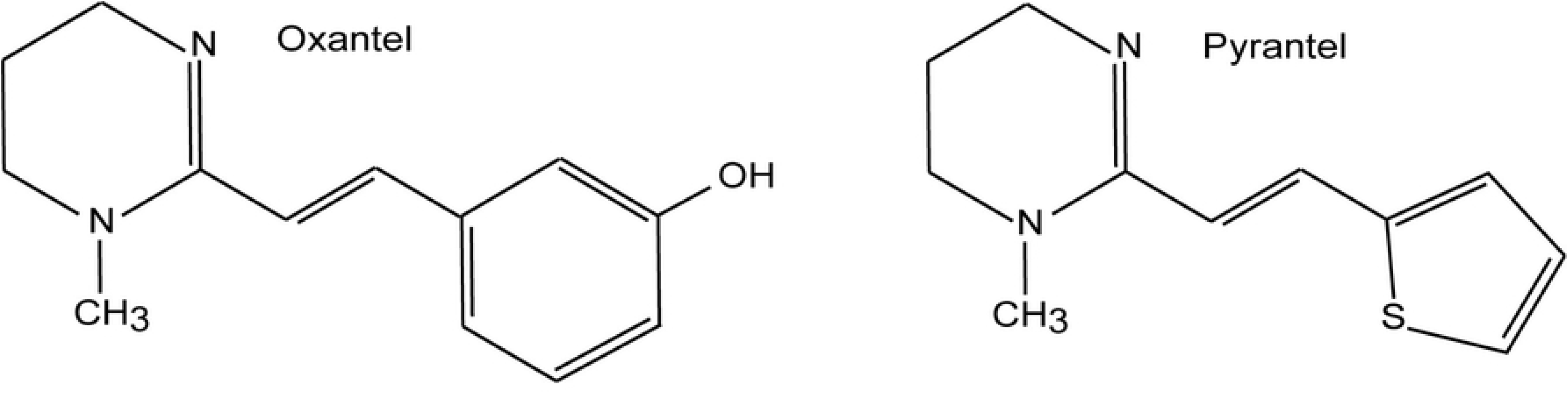
Chemical structure of oxantel and pyrantel. Oxantel: free drawing after https://pubchem.ncbi.nlm.nih.gov/compound/oxantel#section=2D-Structure. Pyrantel: free drawing after https://pubchem.ncbi.nlm.nih.gov/compound/pyrantel#section=2D-Structure

**Figure 5b.**
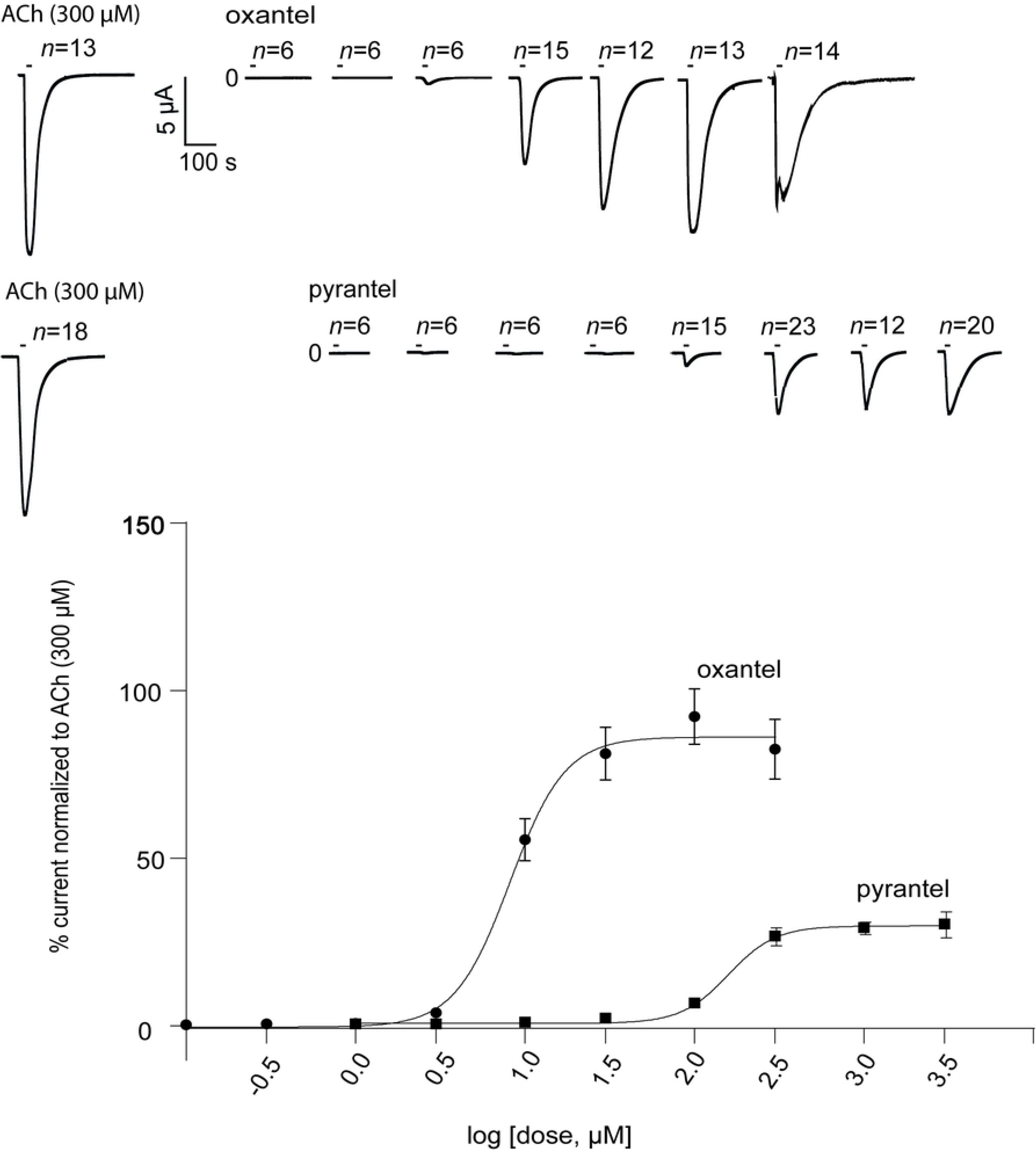
Dose-response curves for oxantel (oxa) and pyrantel (pyr). The current response on *Tsu-*ACR-16-like receptor is normalised to current responses induced by 300 μM ACh and given as mean ± SEM. The EC_50_ ± SD values were 9.48 ± 1.15 μM for oxa and 152.7 ± 1.20 μM for pyr, the relative maximum current responses, *I*_max_ were 86.85 ± 4.63% and 29.41 ± 1.95%, and the Hill slope, *n*_H_, 2.51 ± 1.30 and 3.13 ± 1.07 for oxa and pyr, respectively.

### 2.5. Antagonists

To further characterize the pharmacology of *Tsu-*ACR-16-like receptor, we tested 3 selected antagonists: dihydro-β-erythroidine (DHβE), α-bungarotoxin (α-BTX) and the anthelmintic, derquantel. Fig. 6 shows the effect of 10 μM DHβE and 10 μM derquantel on the *Tsu-*ACR-16-like receptor along with representative current responses. The initial ACh (100 μM) current response of each oocyte was used for normalization, to measure the reduced current responses in the presence of the antagonists. For DHβE, the mean ± SEM inhibition was very small (i.e. 7.60 ± 1.6%) and no inhibition was observed for derquantel (0.16 ± 1.6%). The effect of α-BTX is given in Figure 7A which shows the response-inhibition of the *Tsu-*ACR-16-like receptor to 100 μM ACh when 10 μM α-BTX is applied 10 s before the second application of 100 μM ACh (test oocytes). Fig. 7B shows the effect of the *Tsu-*ACR-16-like receptor when exposed to 100 μM ACh, only (control oocytes). The first current response of 100 μM ACh (ACh1) was used for normalization. Representative current traces along with a bar chart presenting the normalized mean ± SEM of the second and third drug application of test- and control oocytes are given in Fig. 7A and 7B. The 10 μM α-BTX significantly (*P* < 0.0001) reduced the current response of ACh to 8.3 ± 2.4% of the test oocytes (Fig. 7A). This reduction was not observed in the control oocytes (Fig. 7B), thus, when α-BTX was applied 10 s before the second application of ACh, the current response was significantly reduced in test oocytes as compared to control oocytes (Fig. 7A and 7B, *P* < 0.0001).

**Figure 6.**
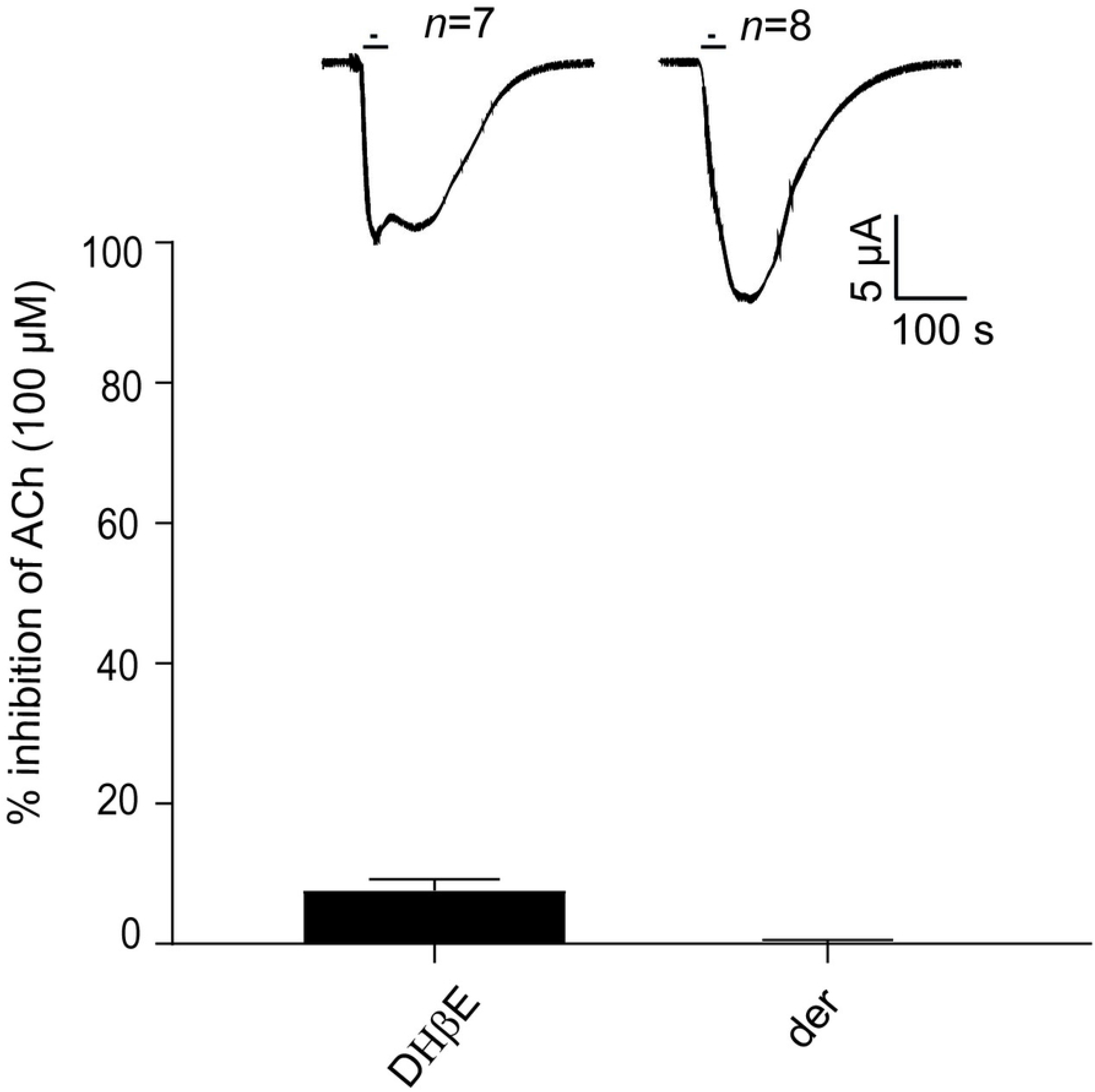
Effect of the antagonists: dihydro-β-erythroidine (DHβE) and derquantel (der) on *Tsu-*ACR-16-like receptor mediated 100 μM ACh current response. Results are given as normalized mean ± SEM inhibition of the initial current response of 100 μM ACh. DHβE produced an almost insignificant block of the *Tsu-*ACR-16-like receptor mediated ACh response (i.e. 7.60 ± 1.61%) and no effect was observed for der (0.16 ± 1.61%).

**Figure 7a and 7b.**
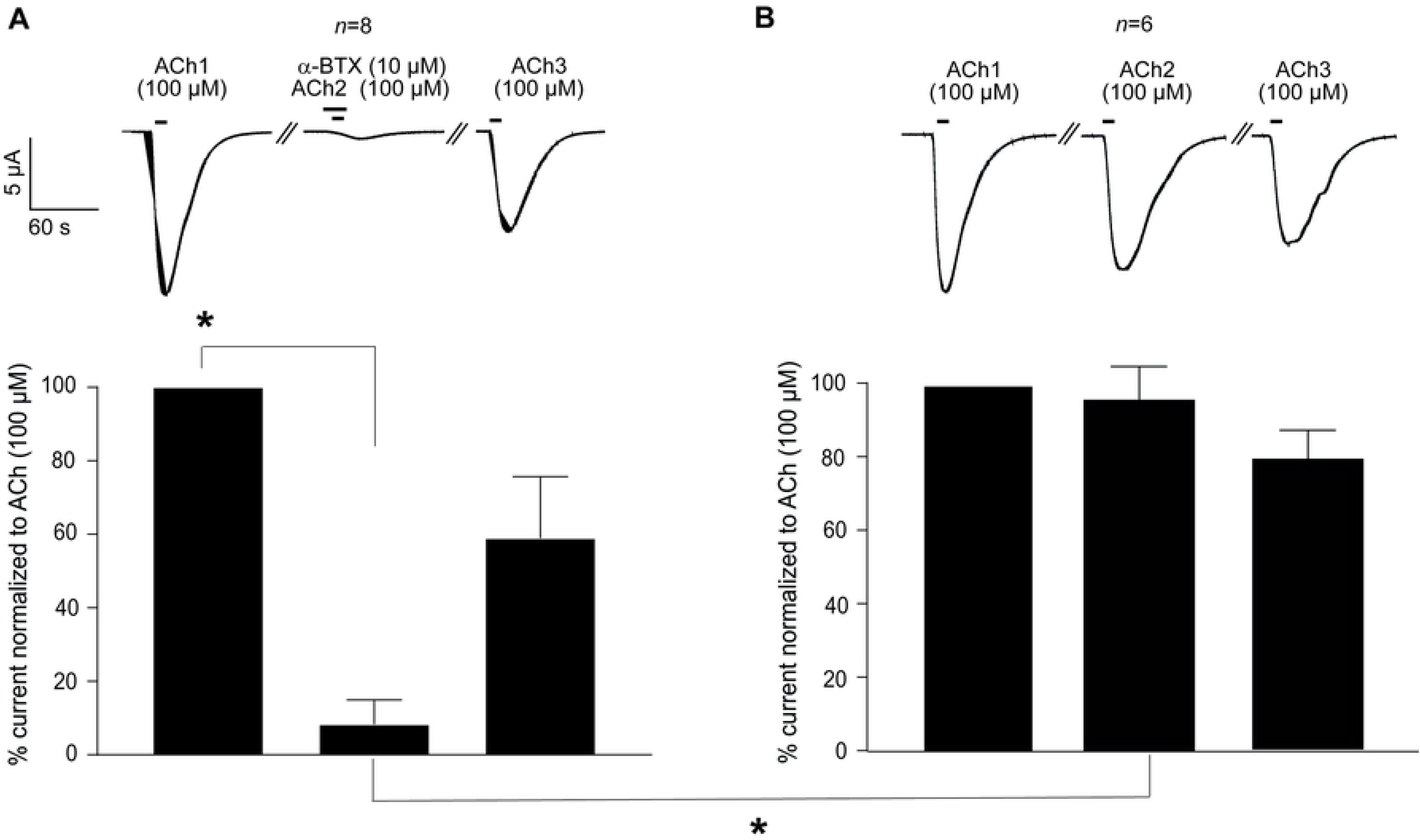
Effect of the antagonists α-bungarotoxin (α-BTX) on *Tsu-*ACR-16-like receptor mediated 100 μM ACh current response. The first ACh current response (ACh1) is set to 100 for both test-(figure 5a) and control oocytes (figure 5b), and subsequent current responses are given as normalized mean ± SEM inhibition of ACh1. *P* < 0.05; significantly different as indicated, Dunnett’s test.

## 3. Discussion

In the present study, we report the identification and the functional expression of the *Tsu-*ACR-16-like receptor, a novel AChR subtype corresponding to the first specific drug target for oxantel to be reported in any nematode species. In reference to the previously reported *L*-AChR, *N*-AChR, and *M*-AChR (respectively for Levamisole-sensitive, Nicotine-sensitive and Morantel- sensitive -AChR subtypes), we named the novel oxantel-sensitive AChR subtype: the *O*-AChR.

### The O-AChR is a novel receptor subtype specific to Clade I nematode species with original pharmacological properties

Use of screens for *C. elegans* mutants that survive exposure to the broad-spectrum anthelmintics provided a means to decipher their molecular targets in a wide range of nematode species. However, this approach was not helpful for oxantel because the *C. elegans* is insensitive to this drug [34]. The weak, but measurable activity of oxantel on recombinant *N*-AChR from *C. elegans* and *A. suum,* supported the hypothesis that a putative ACR-16 homologs in *Trichuris* species could be involved in an oxantel-sensitive receptor. Williamson et al. [35] reported that only two members from the ACR-16 group could be identified in the genomic data from the Clade I species *T. spiralis*. In agreement with this finding, our search for ACR-16 homologs only retrieved two sequences in each of the *Trichuris* species investigated in the present work. The first one corresponded to the highly conserved AChR subunit encoded by the *acr-19* gene; the second one corresponded to a highly divergent subunit specific to Clade I nematode species designated as ACR-16-like.

When co-expressed in the *X. laevis* oocytes with the ancillary protein RIC-3, the ACR-16-like subunits from *T. suis* formed a functional homomeric channel (*O*-AChR) with unexpected pharmacological properties. Indeed, this receptor was highly sensitive to oxantel which is in contrast to the *Asu-*ACR-16 for which a low agonist effect of oxantel has been reported (i.e. <10% of the control ACh current) [29] and to the *Cel-*ACR-16 on which oxantel has an antagonistic effect [30]. Likewise, pyrantel had a relatively high effect on the *Tsu-O*-AChR, whereas pyrantel had no agonist effect on *Asu-*ACR-16 [29] or *Cel-*ACR-16, but in contrast, showed an antagonistic effect on the latter, which was ascribed to pyrantel acting as an open channel blocker [28]. In accordance with this assumption, patch-clamp recording studies from isolated *A. suum* muscle cells, show that both oxantel and pyrantel to act as agonists and open channel blockers [36,37]. Another surprising difference between the *Tsu-O*-AChR and the ACR-16 receptors from *A. suum*, *C*. *elegans* and *P. equorum* is the lack of sensitivity to nicotinic agonists [28,29,33]. Since oxantel has been characterised as an agonist selective for the *N*-subtypes of the ionotropic AChRs [16] which include the ACR-16 receptors, we expected the *Tsu-O*-AChR receptor to be highly sensitive to nicotine, cytosine, 3-bromocytosine, epibatidine and DMPP, but only small current responses were observed using these agonists. Another feature of the *Tsu-O*-AChR receptor is its slow desensitization kinetics, which contrasts with the faster desensitization of the *N*-AChR from *C. elegans* [38], *A. suum* [29] and *P. equorum* [33].

Interestingly, we also showed that the antagonist, α-BTX had a potent inhibitory effect on the ACh induced current responses of the *Tsu-O*-AChR whereas *A. suum* [29] and *C. elegans N*-AChR [28] are nearly insensitive to α-BTX. We point out however, that α-BTX only induced a strong inhibitory effect when α-BTX was applied 10 s before the application of ACh suggesting a slow association time of α-BTX. The *Tsu-O*-AChR was virtually insensitive to DHβE and insensitive to derquantel. This also contrasts with the *Asu*-*N*-AChR which is moderately sensitive to DHβE (~ 65% inhibition) and derquantel (~ 60% inhibition) [29] and the *Cel*-*N*-AChR which is highly sensitive to DHβE [28]. Another notifiable difference is the large current response of the *Tsu-O*-AChR. In the RIC-3 experiments we found a mean current response to 100 μM ACh of 7.54 ± 0.74, μA (*n*=32), whereas the current response of *Asu-N*-AChR to 100 μM ACh was 290.6 ± 63.7 nA (*n*=23) [29], and for *Cel-* ACR-16 to 300 μM ACh ~ 400 nA. For comparison, the current responses to 300 μM ACh of the *Tsu-O*-AChR was 12.94 ± 0.77 μA, *n*=24 (mean ± SEM). Taking into consideration variation between studies, (i.e. number of days for receptor expression), the *Tsu-O*-AChR induced a current response approximately 26 and 19 times higher than *Asu*-*N*-AChR and *Cel*-*N*-AChR, respectively.

Taken together, these results strongly support our hypothesis that *O*-AChR and *N*-AChR represent two distinct class of ionotropic AChR. Despite the important differences, it is noteworthy that there are also similarities between the *O*- and *N*-AChR: both subtypes are not activated by the anthelmintic drugs levamisole or morantel [27–29].

### Sensitivity to oxantel and pyrantel

Small changes in structure of acetylcholine agonists can have large effects on the selectivity and affinity of nicotinic agonists [39]. Perhaps it is not surprising that pyrantel which was modified by replacing the 2-thiophene moiety with the an *m*-oxyphenol group [17] to produce oxantel has a different pharmacology. Thus, the anthelmintic spectrum of oxantel and pyrantel is very different. Pyrantel is a broad-spectrum anthelmintic with no effect on adult *Trichuris* [17] whereas oxantel is a narrow-spectrum anthelmintic with a potent and selective effect on adult *Trichuris* [19–22]. This spectrum difference has previously raised the question whether a cholinergic receptor subtype present in *Trichuris* spp. are different from other intestinal nematode parasites [16], which indeed is now strongly supported by our results.

In conclusion, the discovery of the *Tsu-O*-AChR provides new insights for the high efficacy and specificity of oxantel on whipworms, and provide us with an example of an anthelmintic, that due to its narrow-spectrum will have a lower impact on non-target nematode species. The advantage of such an anthelmintic is the reduced risk of inducing anthelmintic resistance in other parasitic nematode species, and a lower impact on the environmental biodiversity after drug expulsion from the host (i.e. primarily animal hosts).

## 4. Material and methods

### 4.1. Ethic statement

The worm material used in this study was obtained during a previous described study [40] performed at the Experimental Animal Unit, University of Copenhagen, Denmark according to the national regulations of the Danish Animal Inspectorate (permission no. 2015-15-0201-00760). The neurologic tissue from *X. laevis* was obtained from one adult female, which was anaesthetized by submersion into a tricaine solution (ethyl 3-aminobenzoate methanesulfonate, 2g/l) and subsequently decapitated. All procedures involving live material were performed according to the national regulations of the Danish National Animal Experiments Inspectorate (permission no. 2015-15-0201-00560).

### 4.2. Drugs

All drugs except 3-bromocytisine, DHβE, α-BTX and derquantel were purchased at Sigma-Aldrich (Copenhagen, DK). 3-bromocytisine, DHβE and α-BTX were obtained from Tocris Bioscience (Abingdon, UK) and derquantel was purchased at Cayman Chemicals (Ann Abor, MI, USA). Stock solutions of drugs were made in either Kulori medium or DMSO (100%) and stored at −20 or 5°C (i.e. ACh) until use. Before use, stock solutions were dissolved in Kulori medium with a maximum final concentration of DMSO of 0.1%.

### 4.3. Bioinformatics and sequence analysis

The *Asu-*ACR-16 (accession number AKR16139) and the *Cel-*ACR-16 (accession number NP505207) were used as queries in database searches for *Trichuris suis* ACR-16 (*Tsu-*ACR-16) and ACR-16s from other Clade I parasitic nematodes (i.e. *Trichuris* spp. and *Trichinella spiralis*) in the protein-protein BLAST (BLASTp) service at the National Center for Biotechnology Information (NCBI) service [41]. *Cel-*ACR-16 and *Cel-*ACR-19 were used in an alignment with the identified putative ACR-16 sequences from Clade I parasitic nematodes. The accession numbers of the sequences used for the alignment are:

***Caenorhabditis elegans*:** ACR-16 <u>NP505207</u>, ACR-19 <u>NP_001129756</u>. ***Trichuris suis*:** putative ACR-16 <u>KFD48832.1</u> putative ACR-19 <u>KFD70086.1.</u> ***Trichuris trichiura***: putative ACR-16 <u>CDW52185</u>; putative ACR-19 <u>CDW53523</u>. ***Trichuris muris***: putative ACR-16 WBGene00290200; putative ACR-19 WBGene00291941. ***Trichuris spiralis***: putative ACR-16 <u>KRY38920.1</u>, putative ACR-19 <u>KRY27533.1</u>.

Signal peptide was predicted using the SignalP 4.1 server [42] and the transmembrane regions were predicted using the TMHMM version 2 server [43]. Deduced amino-acid sequences were aligned using MUSCLE Phylogenetic analysis was performed on deduced amino-acid sequence. Maximal likelihood phylogeny reconstruction was performed using PhyML V20120412 (https://github.com/stephaneguindon/phyml-downloads/releases) and the significance of internal tree branches was estimated using bootstrap resampling of the dataset 100 times. The accession numbers sequences used for the analysis are:

***Caenorhabditis elegans*:** ACR-7 NP_495647; ACR-8 NP_509745; ACR-9 NP_510285; ACR-10 NP_508692; ACR-11 NP_491906; ACR-12 NP_510262; ACR-14 NP_495716; ACR-15 NP_505206; ACR-16 NP_505207; ACR-19 NP_001129756; EAT-2 NP_496959; LEV-1 NP_001255705; UNC-29 NP_492399; UNC-38 NP_491472; UNC-63 NP_491533.

***Haemonchus contortus***: ACR-16 MH806893. ***Soboliphyme baturini***: acr-16-like VDP07835.1.

***Steinernema glaseri***: ACR-16 KN173365.1. ***Steinernema feltiae***: ACR-19 KN166031.1.

***Toxocara canis***: ACR-16 VDM44142.1; ACR-19 VDM36763.1. ***Parascaris equorum***: ACR-26 KP756902; ACR-27 KP756903. ***Trichuris suis***: ACR-16-like KFD48832; ACR-19 KFD70086‥

***Trichuris muris***: ACR-16-like WBGene00290200; ACR-19 WBGene00291941. ***Trichuris trichiura***: ACR-16-like CDW52185; ACR-19 CDW53523. ***Trichinella spiralis:*** ACR-16-like KRY38920.1; ACR-19 KRY27533.

### 4.4. cDNA synthesis

Worm material was kept in RNA later (Sigma-Aldrich, Copenhagen, DK) at −20°C until use. Total RNA was extracted from whole adult *T. suis* males and females using TRIzol LS Reagent (Invitrogen™). For each isolation, 15 worms were used. The RNA was DNAse treated using DNase I, Amplification Grade (Invitrogen). The whole brain of one *X. laevis* frog was used to extract RNA using Tri™ Reagent®; (MRC. Inc; US) and M tubes in an OctoMacs homogenizer machine (Milteny, Germany) following the manufacturer’ protocol. The isolated RNA was DNAse treated using the RNeasy MinElute Cleanup kit (Qiagen, Germany). The quantity and quality of RNA from *T. suis* and *X. laevis* was assessed by OD measurement in a Nanodrop spectrophotometer (Thermo Scientific, Demark) and by visual inspection in an agarose gel (1%). First strand cDNA was synthesised from 2.5 μg of total RNA from *T. suis* and brain RNA from *X. laevis* using SuperScript IV VILO Master Mix (Invitrogen™) according to manufacturer’s protocol.

### 4.5. PCR and cloning of a full-length T. suis ACR-16 subunit and Xle-RIC-3

To amplify the full-length coding sequence of the *Tsu-acr-16-like* subunit, a specific primer pair, containing the *Bam*HI and the *Not*I restriction enzyme sites and 5 additional nucleotides (*Tsu*-*acr-16*-F-5’-GAATC*-Bam*HI-ATGCGGCCGATAATTTTCCTC-3’ and *Tsu-acr-16*-R-5’-ACGTT *Not*I TCACACAGTTAAATGGGGAGAAC-3’) were designed using the putative *Tsu-acr-16-like* sequence in Wormbase under gene number M514_10316. The specific primer pair used to amplify the *Xle-ric-*3 sequence are described elsewhere [32]. Restriction enzyme sites (*Bam*HI and *Not*I) were included to facilitate ligation into the expression vector pXOOM [44] which was linearized with the restriction enzymes *Bam*HI and *Not*I. PCR amplifications were performed with Platinum SuperFi Green PCR Master Mix (Invitrogen) following the manufacturer’ recommendations. Amplicons were evaluated by gel electrophoresis, digested with *Bam*HI and *Not*I, purified with QIAquick Gel Extraction Kit (Qiagen), cloned into pXOOM and sequenced. A positive clone of *Tsu-acr-16-like* subunit and *Xle-ric-3* was selected and linearized with the restriction enzyme *Nhe*I before *in vitro* transcription using the mMessage mMachine T7 Transcription Kit (Ambion). The cRNA was purified using MEGAclear (Thermo Scientific, Demark). Quantity and quality of cRNA was evaluated by OD measurement in a Nanodrop spectrophotometer (Thermo Scientific, Demark) and by visual inspection in an agarose gel (1%).

### 4.6. Microinjection of Xenopus laevis oocytes

*Xenopus laevis* oocytes were obtained from EcoCyte Bioscience (Castrop-Rauxel, Germany) and kept at 19°C in Kulori medium (90 mM NaCl, 4 mM KCl, 1 mM MgCl_2_, 1 mM CaCl_2_, 5 mM HEPES, pH:7.4) until injection. Oocytes were injected with 50 nl of cRNA in RNAse-free water using a microinjector (Nanojet, Drummond Broomal, PA, USA). To test *ric*-3 effects on the receptor expression, 25 ng *Tsu-acr-16-like* cRNA was injected alone or with either 5 ng *Asu*-*ric*-3 or 5 ng *Xle*-*ric*-3 cRNA. To exclude endogenous nAChR expression induced by *Asu*-*ric*-3, 5 ng *Asu*-*ric*-3 was injected alone. These amounts of cRNA were chosen in order to compare the drug-potency results of the *Tsu-*ACR-16-like receptor with that of *Asu-*ACR-16[29]. Only half (i.e. 12.5 ng *Tsu-acr-16-like* cRNA and 2.5 ng *Asu*-*ric*-3) was used for the dose-response curves and antagonist analysis. To allow for receptor expression, the injected oocytes were incubated in Kulori medium at 19°C for 3-7 days and the Kulori medium was changed daily.

### 4.7. Two-electrode voltage clamp of Xenopus laevis oocytes

Two-electrode voltage-clamp (TEVC) recordings were obtained using an Oocyte Clamp Amplifier OC-725 B (Warner Instruments Corp., USA) connected to an Axon Digidata^®^ 1440A digitizer (Axon Instruments, Molecular Devices, USA) and was performed at ~19°C under continuous flow of Kulori medium with the oocytes clamped at −60 mV. Data were sampled at 2 kHz using the pClamp 10.4 acquisition software (Axon Instruments, Molecular Devices, USA). The microelectrodes were pulled from glass capillaries (TW 120.3, World precision instruments, USA) on a programmable micropipette puller (Narishige, Japan). The resistance when filled with 3 M KCl ranged from 0.5 to 1.5 MΩ. Ag/AgCl reference electrodes were connected to the bath with agar bridges. For a minimum of 4 hours prior to recording, all oocytes were incubated in BAPTA-AM at a final concentration of 100 μM to chelate intracellular Ca^2+^ ions and hereby prevent activation of endogenous calcium activated chloride channels during recordings. Recordings from non-injected oocytes were used as negative controls.

### 4.8. Drug-potency-tests

All agonists were used at a final concentration of 100 μM and were tested in 3-4 experiments using 3-4 different batches of oocytes batches. For each experiment, 6 oocytes were tested per drug. For each experiment, 6 oocytes were exposed to 100 μM ACh for 10 s, and all other responses in the same experiment, were normalized to the mean response of these controls. Initial experiments showed a consistent decrease in the response to 100 μM ACh when applied repeatedly, even after washing periods for up to 5 min between agonist applications. Therefore, each drug was tested on oocytes not previously exposed to ACh (100 μM). The total number of oocytes examined per drug was: *n* = 23 for oxantel, *n* = 16 for pyrantel, *n* = 15 for epibatidine, *n* = 16 for nicotine, *n* = 15 for 3- bromocytisine, *n* = 16 for DMPP, *n* = 17 for morantel, *n* = 17 for cytisine and *n* = 15 for levamisole. Each drug was applied for 10 s followed by wash off until the current had returned to pre-stimulation values.

### 4.9. Dose-response studies

The dose-response studies were in total performed on 3 different oocyte batches. For oxantel the number of measurements per drug concentration were as follows: 0.3 μM, *n*=6; 0.1 μM, *n*=6; 3 μM, *n*=6; 3 μM, *n*=6; 10 μM, *n*=15; 30 μM, *n*=12; 100 μM, *n*=13; 300 μM, *n*=14. For pyrantel *n* per drug concentration were: 1 μM, *n*=6; 3 μM, *n*=6; 10 μM, *n*=6; 30 μM, *n*=6; 100 μM, *n*=15; 300 μM, *n*=23; 1000 μM, *n*=12; 3000 μM, *n*=20. For each experiment, 6 oocytes were exposed to 300 μM ACh for 10 s, and all other responses in the same experiment, were normalized to the mean response of these positive controls. The ACh concentration of 300 μM was used to reach the supposed maximal activation of the *Tsu-*ACR-16-like receptor. Each drug and drug concentration were tested as described for the drug-potency-tests.

### 4.10. Antagonists

The effects of the antagonists DHβE, derquantel and α-BTX (10 μM) were examined in the presence 100 μM ACh as previously described for *Asu-*ACR-16 [29]. In short, *X. laevis* oocytes co-injected with *Tsu-acr-16-like-* and *Xle-ric*-3 cRNA were sequentially superfused with ACh for 10 s, then ACh + antagonist for 10 s, and finally with ACh for 10 s. For α-BTX a five-step protocol including a pre-incubation (10 s) with the antagonist (10 μM) was used with ACh (100 μM). *Xenopus laevis* oocytes, co-injected with *Tsu-acr-16-like-* and *Xle-ric*-3 cRNAs, were exposed to: i) a control application of 100 μM ACh for 10 s (first application); ii) followed by a wash-off period of 5 min; iii) then by an application of 10 μM α-BTX for 10 s, immediately followed by 100 μM ACh and the continued presence of α-BTX for 10 s (second application); iv) then a wash-off period of 5 min; v) and finally an application of ACh for 10 s (third application). Control oocytes were exposed to ACh for 10 s in 3 consecutive steps, each separated by a wash-off period of 5 min. For each antagonist, *n*=6-8.

### 4.11. Electrophysiological data and statistically analysis

All acquired electrophysiological data were analysed with Clampfit 10.7 (Molecular Devices, Sunnyvale, CA, USA) and GraphPad Prism 8 (GraphPad Software, La Jolla, CA, USA) and from all experiments, peak currents from BAPTA-AM-incubated oocytes were measured after application of drugs. For the auxiliary protein (RIC-3) test, the largest group mean current in response to 100 μM ACh was set to 100%, and all other responses were normalized to this. The relative means were statistical analysed using One-Way ANOVA with a Dunnett’s Test and *P* < 0.05 was considered significant. For the drug-potency-test, peak currents of drugs were normalized to the peak current measured in the presence of 100 μM ACh and was expressed as mean ± SEM. Data was tested for normality using the D’Agostino-Pearson normality test. Drug-group means were statistical analysed using One-Way ANOVA with a Turkey’s Multiple Comparison Test where *P* < 0.05 was considered significant.

For the dose-response relationships, and for each experiment, 6 oocytes were exposed to ACh (300 μM) for 10 s, and all other drug responses in the same experiment, were normalized to the mean response of these controls. The normalized current as a function of drug concentration allowed fitting the dose-response curves with a Hill equation, using nonlinear regression analysis with a variable slope model in GraphPad Prism 8. The following equation was used:

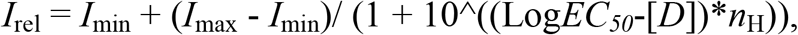

where *I*_rel_ is the mean relative current, *I*_max_, is the relative current obtained at saturating agonist concentration, *I*_min_ is the relative current obtained at agonist concentration 0 μM, *EC*_*50*_ is the concentration of agonist resulting in 50% of the maximal current response, [*D*] is the drug concentration and *n*_H_ is the Hill coefficient. *I*_max_, *EC*_*50*_ and *n*_H_ were fitted as free parameters whereas *I*_min_, was constrained to 0.

For the antagonist test with α-BTX, the current response of α-BTX and ACh in the continued presence of α-BTX (second application) was normalized to the first response at 100 μM ACh (first application) which was set to 1. The group mean of the α-BTX response were statistical analysed using One-Way ANOVA with a Dunnett’s test.

## 6. Acknowledgement

The study was supported by the Independent Research Fund Denmark (DFF – 4184-00210), the Danish National Advanced Technology Foundation (5184-00048B) and the Lundbeck Foundation (R9-A1131). authors would like to thank Richard Martin at Iowa State University, USA for providing the *Asu-*ACR-16 and *Asu-*RIC-3. RJM is supported by NIH, the National Institute of Allergy and Infectious Diseases grants R01AI047194-17, R21AI092185-01A1 and the E. A. Benbrook Foundation for Pathology and Parasitology. In addition, the authors wish to thank Vibeke Grøsfjeld Christensen at the Veterinary and Animal Sciences, University of Copenhagen for technical assistance.

## Supplementary figure legends

**Supplementary figure 1**

**Distance tree showing relationships of the ACR-16-like acetylcholine receptor (AChR) subunits from Clade I nematode species, with other AChR subunits from the ACR-16 group from nematode species representative from Clade III, Clade IV, and Clade V.**

NJ-Tree was built upon an alignment of AChR subunit deduced amino-acid sequences. The tree was rooted with the *C. elegans* UNC-63 sequence. Scale bar represents the number of substitutions per site. Boostrap values (1000 replicates) are indicated on branches. Accession numbers for sequences used in the analysis are provided in Material and Methods section. Nematode clades refer to Blaxter et al. 1998 [1]. AChR subunit sequences from Clade I species are highlighted in red, AChR subunit sequences from Clade III species are highlighted in blue, AChR subunit sequences from Clade V species are highlighted in pink (in black for *C. elegans*). *Cel*, *Hco*, *Sba*, *Sgl*, *Sfe*, *Tca*, *Tsp*, *Tsu*, *Ttr* and *Tmu* refer to: *Caenorhabitis elegans*, *Haemonchus contortus*, *Soboliphyme baturini*, *Steinernema glaseri*, *Steinernema feltiae*, *Toxocara canis*, *Trichinella spiralis*, *Trichuris suis*, *Trichuris trichiura*, and *Trichuris muris* respectively.

**Supplementary figure 2**

**Desensitization kinetics of *Tsu-*ACR16-like receptor.** A representative response of the *Tsu-* ACR16-like receptor to 1 min exposure of 100 μM ACh. The *Tsu-*ACR16-like receptor is characterized by a slow-desensitization kinetic as compared to *Asu*-ACR-16 [29] and *Peq*-ACR-16 [33].

